# Assembly principles of a SYCP2-HORMAD1-HORMAD2 mammalian meiotic axis complex

**DOI:** 10.64898/2026.06.03.730022

**Authors:** Elizaveta Selezneva, Franziska Müller, Petra Janning, John R. Weir

**Affiliations:** Friedrich Miescher Laboratory of the Max Planck Society, Max-Planck-Ring 9, 72076, Tübingen, Germany; Max Planck Institute for Molecular Physiology, Otto-Hahn-Str. 11, 44227 Dortmund, Germany

## Abstract

During meiotic prophase I, the chromosome axis orchestrates programmed DNA double-strand break formation, repair and synapsis between homologous chromosomes. In mammals, the axis is assembled from the coiled-coil elements SYCP2 and SYCP3 that come together with the HORMA-domain proteins HORMAD1 and HORMAD2, but how these components associate into a coherent structural unit remains incompletely understood. Combining recombinant reconstitution, mass photometry, SEC-MALS, AlphaFold modelling and crosslinking mass spectrometry, we show that the HORMA domains of HORMAD1 and HORMAD2 form a selective pseudosymmetric heterodimer independently of either protein’s own closure motif, with interface determinants conserved across vertebrates. We identify a previously unrecognised second closure motif (CM2) in SYCP2 that preferentially binds HORMAD1, distinct from the previously described HORMAD2-binding closure motif (CM1). Together, these results revise the current model of mammalian axis assembly and define a tripartite SYCP2-HORMAD1-HORMAD2 module as a fundamental structural unit of the mammalian meiotic chromosome axis.

## Introduction

Proper segregation of homologous chromosomes during meiosis is ensured by crossovers, which are derived from programmed double-strand breaks (DSBs) during prophase I (reviewed in (Zickler & Kleckner, 2023)). Synapsis between homologous chromosomes facilitates the proper formation of crossovers and also regulates their spatial distribution. DSBs are, on the one hand, necessary for the proper physical linkage between homologs, but on the other hand, carry potential risks for genome integrity; therefore, their number, localisation, and timing must be strictly regulated. DSBs are formed by the type II topoisomerase complex SPO11-TOPOVIBL (Bergerat et al., 1997; Keeney et al., 1997; Robert et al., 2016) and recruited to the chromosome axis by several auxiliary proteins, REC114, MEI4 and IHO1 (Rec114, Mei4 and Mer2 in budding yeast) (Kumar et al., 2010, 2018; Panizza et al., 2011; Stanzione et al., 2016).

In prophase I, sister chromatids are organised into linear arrays of chromatin loops anchored to a proteinaceous axis (Novak et al., 2008; Zickler & Kleckner, 2023). Loop organisation is established by the meiosis-specific cohesin complex (SMC1β, SMC3, STAG3, and the kleisin REC8 or RAD21L; (Grey & de Massy, 2021)), while the axis itself is built from two structurally distinct conserved classes of protein: long coiled-coil “axial elements” - SYCP2 and SYCP3 in mammals, Red1 in budding yeast - and HORMA-domain proteins - HORMAD1 and HORMAD2 in mammals, Hop1 in yeast. The HORMA fold, named for its three founding members (Hop1, Rev7 and Mad2) (Aravind & Koonin, 1998), comprises an N-terminal globular core followed by a topologically mobile “safety-belt” element (β7-β8) that can rearrange to encircle a short peptide closure motif (CM) present either on an interacting partner or on the HORMA protein’s own C-terminus (Sironi et al., 2002). Engagement of the safety-belt over a closure motif locks the HORMA domain into a “closed” topology, and the open-to-closed transition is remodelled by the AAA+ ATPase TRIP13 (Pch2 in yeast), which engages the HORMA N-terminus to release the safety-belt from its bound partner (Figure 1B) (Gu et al., 2022; Vader, 2015; Ye et al., 2017)

**Figure 1.**
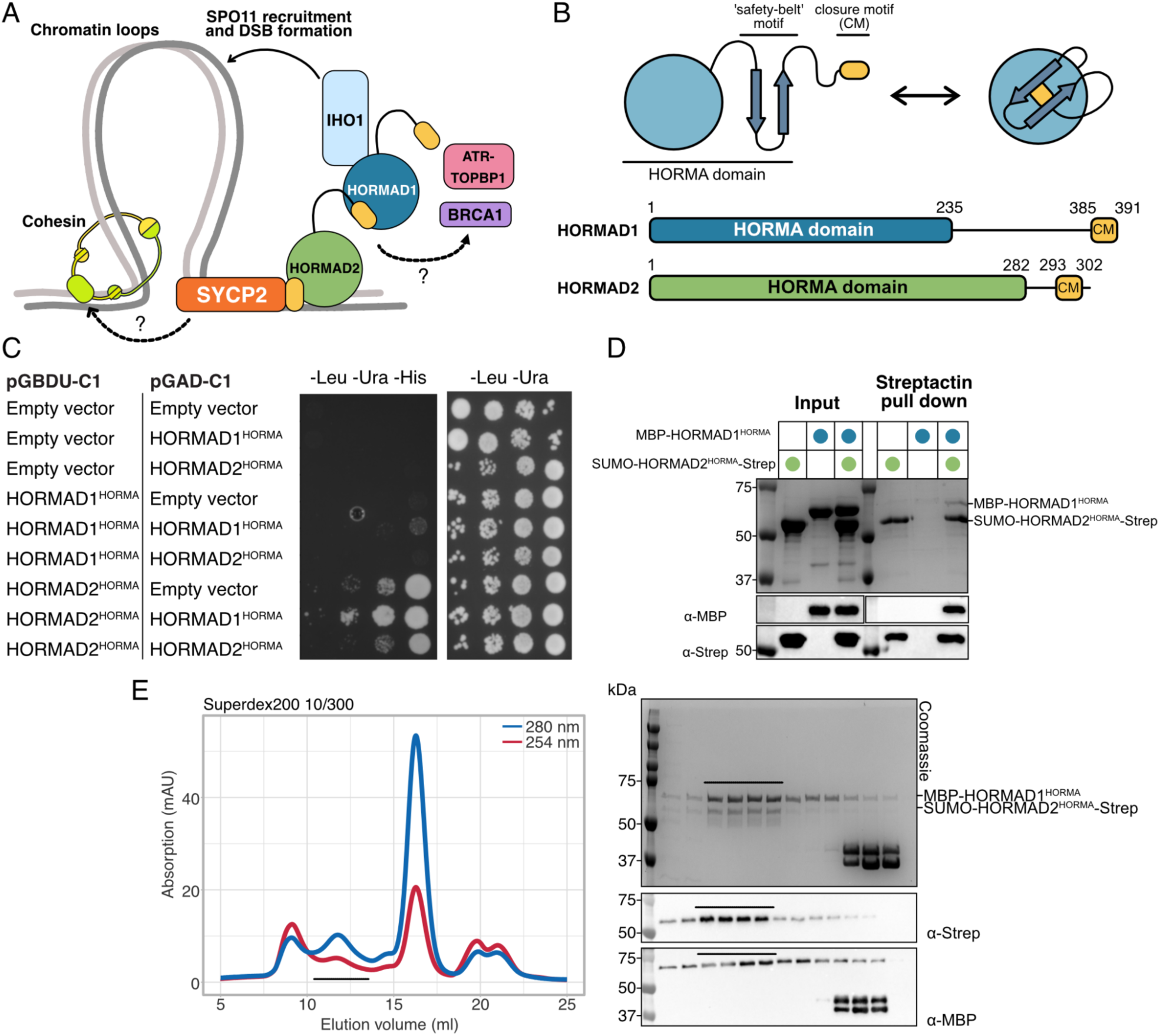
HORMAD1 and HORMAD2 form a complex. A) Introductory cartoon. Sister chromatids (grey and black lines) are entrapped by cohesin. Cohesin also produced loops of chromatin. SYCP2 (orange) might interact directly with meiotic cohesin. SYCP2 contains a closure motif (yellow) that binds to HORMAD2 (green). HORMAD2 contains a closure motif (yellow) that can be bound by HORMAD1 (blue). Both HORMADS are involved in the recruitment of ATR-TOPBP1 and BRCA1 by unknown mechanisms. HORMAD1 binds directly to IHO1, which in turn recruits and activates the SPO11 machinery to make double-stranded DNA breaks and initiate meiotic recombination. B) HORMAD1 and HORMAD2 both consist of an N-terminal HORMA domain which can bind in cis to a C-terminal closure motif, locking the HORMA domain in a “closed” topological conformation. C) Yeast two-hybrid experiments with either HORMAD1, HORMAD2 or an empty vector used as indicated. The drop assay was performed with serial dilution 1:1000, 1:100, 1:10 and undiluted sample with OD600 0.5 D) Streptactin pull down. Combinations of MBP-HORMAD1^HORMA^ and Strep-HORMAD2^HORMA^ were incubated overnight and then incubated for 30 minutes on Streptactin beads before the beads were washed and eluted. Both protein identity and a lack of non-specific binding were confirmed using western blotting. E) Left, chromatogram of size exclusion chromatography (SEC) experiment on Superdex 200 10/300 of co-expressed and affinity-purified MBP-HORMAD1^HORMA^/SUMO-HORMAD2^HORMA^-Strep complex; right Coomassie stained SDS-PAGE and Western blots from the SEC.

In mammals, HORMAD1 and HORMAD2 are loaded onto unsynapsed axes at the onset of prophase I (Fukuda et al., 2010; Wojtasz et al., 2009) and are essential for the proper initiation of meiotic recombination. HORMAD1 recruits the DSB-promoting machinery to the axis through a direct interaction with the C-terminus of IHO1, a component of the IHO1-MEI4-REC114 complex (orthologous to the Mer2-Mei4-Rec114, or “RMM”, complex of *S. cerevisiae*) that licenses SPO11-TOPOVIBL activity (Biot et al., 2024; Dereli et al., 2024; Laroussi et al., 2023; Stanzione et al., 2016). Consistent with this, *Hormad1*-deficient mice show severely reduced DSB formation, impaired homolog pairing, and infertility in both sexes (Daniel et al., 2011; Shin et al., 2010). HORMAD1 is also required to bias DSB repair towards the homolog rather than the sister chromatid, a function conserved with budding yeast Hop1 (Carballo et al., 2008; Carofiglio et al., 2018; Shin et al., 2013).

Beyond their roles in DSB formation, HORMAD1 and HORMAD2 are the upstream sensors of the meiotic prophase surveillance system that eliminates meiocytes carrying persistent asynapsis or unrepaired DSBs (reviewed in (Subramanian & Hochwagen, 2014). Both HORMADs accumulate on unsynapsed axes, where they recruit and locally concentrate the DNA-damage-response kinase ATR (Wojtasz et al., 2012). Failure of synapsis or DSB repair triggers oocyte elimination perinatally through a network that involves CHK2 and BRCA1 (Bolcun-Filas et al., 2014; Ravindranathan et al., 2022); recent work has suggested that BRCA1 is recruited to unsynapsed axes through an interaction with the HORMA domain of HORMAD1 (Bai et al., 2024; Jiao et al., 2025) (Figure 1A). The correct stoichiometric assembly, axial recruitment, and synapsis-coupled removal of HORMAD1 and HORMAD2 are therefore central not only to the structural organisation of the axis but to the regulation of recombination and gamete viability.

Despite this central role, the molecular basis for HORMAD recruitment to the mammalian axis is incompletely understood. SYCP2 comprises an N-terminal ARM-like region and a pleckstrin-homology (PH) domain (Feng et al., 2017), a long predicted intrinsically disordered central region, and a C-terminal coiled-coil that forms transverse filaments with SYCP3 (Syrjänen et al., 2014; West et al., 2019). A closure motif within the disordered region of SYCP2 binds the HORMA domain of HORMAD2 (West et al., 2019), and a co-complex of HORMAD1 bound to the closure motif of HORMAD2 has subsequently been crystallised (Wang et al., 2023). In a recent *in vivo* test of this model, targeted disruption of the SYCP2 closure motif in mice selectively abolishes HORMAD2 - but not HORMAD1 - axial localisation, with phenotypes indistinguishable from *Hormad2*-null animals, establishing the SYCP2-HORMAD2 interaction as the essential scaffold for synapsis surveillance and implying that HORMAD1 is delivered to the axis by an additional, undefined route (Raveendran et al., 2026). The same study, using yeast two-hybrid assays, further suggests that the HORMA domains of HORMAD1 and HORMAD2 themselves interact, independently of either protein’s closure motif (Raveendran et al., 2026). In parallel, we have shown in budding yeast that the HORMA domain of Hop1 engages the ARM-PH region of Red1, defining a second, closure-motif-independent mode of HORMA recognition by an axial element (Chen et al., 2025). How a single SYCP2 molecule accommodates two functionally distinct HORMADs with apparently divergent recruitment requirements, whether an analogous HORMA-axial-element interaction operates in mammals, and whether the proposed HORMA-HORMA heterodimer is biochemically robust and structurally defined therefore all remain open questions.

Here we address these questions by reconstituting and characterising a mammalian SYCP2-HORMAD1-HORMAD2 complex. We show that the HORMA domains of HORMAD1 and HORMAD2 form a closure-motif-independent heterodimer with defined molecular determinants of selectivity that are conserved across vertebrates. We identify a previously unrecognised second closure motif (CM2) within SYCP2 that preferentially engages HORMAD1, distinct from the HORMAD2-binding CM1 - providing a structural explanation for the selective HORMAD1 axial recruitment observed in SYCP2-CM1-deficient mice. We further demonstrate that HORMAD1 can engage the ARM-PH region of SYCP2 in the absence of HORMAD2, paralleling our recent findings in yeast, and that this binding mode is suppressed when HORMAD2 is present. Together these results revise the prevailing model of mammalian axis assembly and define a tripartite SYCP2-HORMAD1-HORMAD2 module as a fundamental structural unit of the mammalian meiotic chromosome axis.

## Results

### HORMAD1 and HORMAD2 form a complex

As described above, recent work has shown that the HORMA domains HORMAD1 and HORMAD2 could interact in the absence of SYCP2 and in the absence of apparent closure motifs, via yeast two-hybrid (Raveendran et al., 2026). We confirmed this with our own yeast two-hybrid experiments. We found indications that the HORMA domains of HORMAD1 (residues 1-235 from hereon HORMAD1^HORMA^) and HORMAD2 (residues 1-282, from hereon HORMAD2^HORMA^) can interact, but HORMAD1^HORMA^ showed no evidence of self-interaction, neither did HORMAD2^HORMA^ (Figure 1C).

We asked whether an interaction between HORMAD1^HORMA^ and HORMAD2^HORMA^ could be reconstituted with recombinant proteins. To test this, we purified HORMAD1^HORMA^ with an N-terminal MBP tag (MBP-HORMAD1^HORMA^), and HORMAD2^HORMA^ with an N-terminal SUMO-tag and a C-terminal 2xStrep-II tag (SUMO-HORMAD2^HORMA^-Strep) (Supplementary Figure 1A and B). Consistent with the Y2H data, we observed complex formation for the separately purified HORMAD1^HORMA^ + HORMAD2^HORMA^ (Figure 1D), though this required an overnight incubation of HORMAD1^HORMA^ and HORMAD2^HORMA^. Using coexpression of two HORMA-domains (N-terminal 6xHis-MBP-tag for HORMAD1^HORMA^ and N-terminal 6xHis-SUMO and C-terminal Strep-tag for HORMAD2^HORMA^), we purified a complex of HORMAD1^HORMA^ and HORMAD2^HORMA^ for further analysis, which appeared to elute as a single species in size exclusion chromatography (Figure 1E).

### Characterisation of a HORMAD1-HORMAD2 complex

Having established that a heterotypic HORMAD1^HORMA^ -HORMAD2^HORMA^ complex can be formed, we asked what the stoichiometry of these complexes was. Utilising mass photometry, we observed a single species of 107 kDa (Figure 2A). This species is consistent with an MBP-HORMAD1^HORMA^/SUMO-HORMAD2^HORMA^-Strep heterodimer (theoretical molecular mass of 119 kDa). Since mass photometry is carried out at low nanomolar concentrations, in this case ∼30 nM, we could not exclude that we observed the disintegration of a larger complex. Therefore, we measured the mass of the HORMAD1^HORMA^/HORMAD2^HORMA^ complex using SEC-MALS (Figure 2B), where we observed a contiguous peak, rather than separate peaks. We determined species corresponding to 61.2 kDa (green trace), 78.4 kDa (yellow trace) and 111.3 kDa (red trace). While there are some inconsistencies with the theoretical and measured molecular masses, we assume that the right hand side of the peak corresponds to HORMAD^HORMA^ monomers, and the left part of the peak to a heterodimer (Figure 2B). Taken together, we conclude that HORMAD1^HORMA^ / HORMAD2^HORMA^ predominantly form a heterodimer in solution.

**Figure 2.**
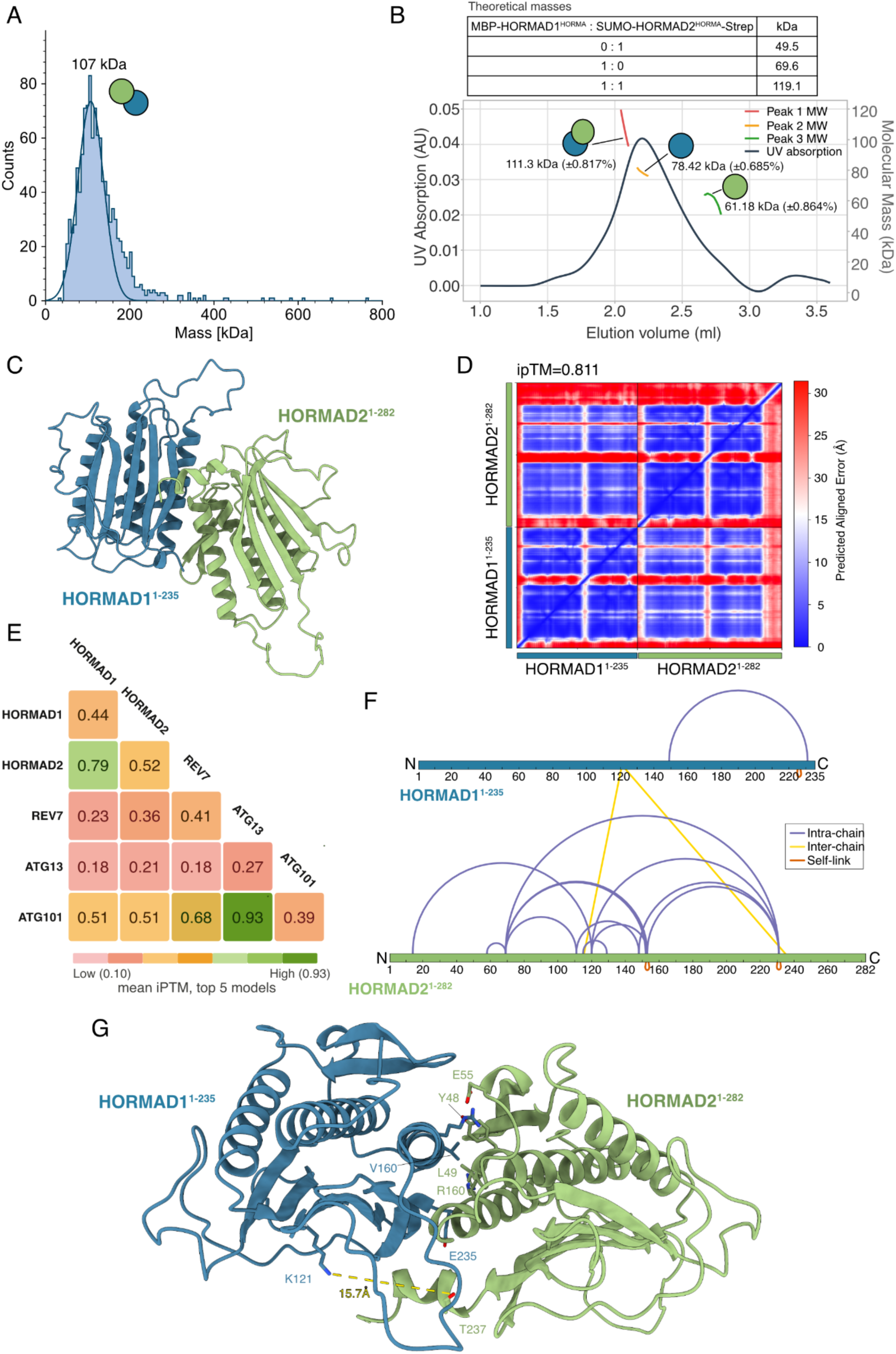
Characterisation of a HORMAD1-HORMAD2 complex. A) Mass photometry profile of purified HORMAD1-HORMAD2 heterodimer at 30 nM, recorded for 60 seconds. B) SEC-MALS of purified coexpressed HORMAD1-HORMAD2 as described in Methods. Sample was run on the column Superdex200 5/150 (Cytiva) in 20 mM HEPES pH 8.2, 150 mM NaCl, 1 mM TCEP at the concentration ∼10 μM. C) Ribbon representation of the HORMAD1^HORMA^-HORMAD2^HORMA^ predicted heterodimer. HORMAD1 is coloured blue, HORMAD2 in green. We have removed the low confidence (pLDDT < 30, see Supplementary Figure 2B) regions (HORMAD1 residues 1-15 and HORMAD2 residues 1-20 and 247-282) for greater clarity. D) PAE plot of the predicted HORMAD1^HORMA^-HORMAD2^HORMA^ heterodimer. E) Summary of ipTM scores for pairwise predictions as shown. 25 predictions were carried out for each pair. The mean ipTM for the top 5 scoring models is shown. Distribution of all 25 ipTM values is shown in Supplementary Figure 2A. F) XL-MS map of high confidence (score cut off to ensure false discovery rate of ≤1%) intra-chain (blue), inter-chain (green) and self-links (orange). G) Alphafold 2 prediction with mapped crosslink between HORMAD1 K121 and HORMAD2 T237; the stick cartoons of the side chains of the residues on potential interface are labelled.

Having established the basic stoichiometry of the HORMAD1^HORMA^/HORMAD2^HORMA^ complex as 1:1, we turned to AlphaFold to build potential models of the heterodimeric complex. We explored both AlphaFold2 and AlphaFold3 predictions, but AlphaFold2 gave consistently higher ipTM scores for the models, and was subsequently used in all predictive approaches described here. Heterodimers of HORMAD1^HORMA^/HORMAD2^HORMA^ were predicted with high confidence (average ipTM 0.79 for the top 5 models of 25 predictions, ipTM of 0.81 for the top model) (Figure 2C and D), whereas, consistent with our experimental observations, homodimeric models were of low confidence (HORMAD1-HORMAD1 = average ipTM 0.44; HORMAD2-HORMAD2 = average ipTM 0.52, Figure 2E, Supplementary Figure 2A). We further explored the possibility of a hallucinated HORMA dimer interface, due to a bias from the AlphaFold2 training data, given the diverse array of HORMA-dimers that have been structurally characterised (reviewed in (Gu et al., 2022)). We explored all possible HORMA dimer combinations for HORMAD1^HORMA^, HORMAD2^HORMA^, REV7, ATG13^HORMA^ and ATG101 all from *Mus musculus* proteome. We found consistently low or borderline ipTM scores for all HORMA pairs except for HORMAD1-HORMAD2 and the *bona fide* ATG13^HORMA^-ATG101 (ipTM 0.93), for which there are crystal structures of the fission yeast (4YK8, (Suzuki et al., 2015) and human orthologs (5C50 (Qi et al., 2015)) (Figure 2E, Supplementary Figure 2A). Analysis of the HORMAD1^HORMA^ / HORMAD2^HORMA^ predicted model showed both HORMA domains to be in the closed conformation (Figure 2C), and the predicted structure of HORMAD1^HORMA^ was essentially identical to the experimental structure of HORMAD1^HORMA^ in complex with HORMAD2 closure motif (PDBID 8J69 (Wang et al., 2023), C-alpha RMSD of 0.6Å over 191 residues).

For further validation of the HORMAD1^HORMA^ / HORMAD2^HORMA^ model, we subjected our protein complex to chemical crosslinking with DSBU, followed by trypsination and mass-spectrometry (XL-MS) (Figure 2F). We then modelled the cross-links onto the HORMAD1^HORMA^ / HORMAD2^HORMA^ AlphaFold prediction. This approach revealed one very high-confidence cross-link between HORMAD1 and HORMAD2 between HORMAD1 K121 and HORMAD2 T237, and a second cross-link of HORMAD1 K121 to K115 of HORMAD2. On the model, this cross-link has an NZ to OG distance of 15Å, which is consistent with the length of the DSBU cross-linker of 12.5Å (Figure 2G). The second cross-link, HORMAD1 K121 to HORMAD2 K115, has a longer NZ to NZ distance of 23Å, K115 is in a flexible loop, which could conceivably move closer to HORMAD1 K121. Taken together, we consider that the HORMAD1^HORMA^ / HORMAD2^HORMA^ prediction shown here is likely an accurate reflection of one conformation of the HORMAD1^HORMA^ / HORMAD2^HORMA^ heterodimer.

### The HORMAD1-HORMAD2 interface is evolutionarily conserved

Meiotic HORMADs are predicted to be ubiquitously conserved across eukaryotes (Tromer et al., 2021), and have formally been shown to be present in green plants (Caryl et al., 2000), brown algae (Kane et al., 2025), budding yeast (Hollingsworth et al., 1990), fission yeast (Lorenz et al., 2004), nematodes (Zetka et al., 1999) and vertebrates (Wojtasz et al., 2009). Most clades have a single meiotic HORMAD gene, though the emergence of at least two copies has occurred multiple times, presumably independently, in evolution. We asked what features differentiate HORMAD1 from HORMAD2, which of these features facilitate the heterodimer interface, and are these features conserved? We compared HORMAD1 and HORMAD2 sequences from 132 vertebrate species. HORMAD1 consistently had a C-terminal extension, with a median length of 87 amino acids, with mouse HORMAD1 having an 86 amino acid extension. Between the conserved HORMA domains, the average sequence identity was 54% and similarity 70% over 284 positions (Supplementary Figure 3).

We next took a closer look at the HORMAD1-HORMAD2 model with particular regard for the interface. We first examined the pLDDT scores. As expected, the termini of the models had low pLDDT scores, corresponding to their lack of predicted structure. We therefore excluded all the residues at the model termini with a pLDDT of <50 (HORMAD1 1-15; HORMAD2 residues 1-20; 247-282) (Supplementary Figure 2B). The HORMA domains are essentially identical to one another by gross morphology. HORMAD2 packs principally against αA and β7 of HORMAD1 via αA, αA1, αC and the β1-β2 hairpin (Figure 2G). Given the very high level of sequence identity within these regions, this interface alone might allow the formation of homo-dimers, but these are clearly disfavoured as shown by predictive and experimental approaches. The prediction suggests that selectivity is provided by two mechanisms. Firstly in the β1-β2 hairpin, S56 and S67 of HORMAD2 are replaced with R52 and C63 in HORMAD1. Secondly, there is a salt bridge between HORMAD1 E235 of αD and HORMAD2 R160 of αC, which is dependent on the varied region of HORMAD1 C-terminal to the HORMA domain core. Based on our sequence alignments, we expect the vast majority of vertebrate HORMAD1 and HORMAD2 HORMA domains will form heterodimers analogously to the mouse proteins. It is also feasible that homodimers may form under certain conditions, since none of predicted side-chain interactions would forbid homodimer formation, rather favour heterodimers.

### HORMAD1 and HORMAD2 bind via distinct closure motifs in SYCP2

SYCP2 contains N-terminal ARM and PH domains (Feng et al., 2017), followed by a closure motif that has been shown to bind directly to HORMAD2 (West et al., 2019) (Figure 3A). We asked via AlphaFold predictions whether the HORMAD1-HORMAD2 heterodimer could still bind to SYCP2 and whether the N-terminal globular domains of SYCP2 might bind to the HORMA domains, as we have recently shown for the meiotic HORMA Hop1 and Red1 in budding yeast (Chen et al., 2025). Surprisingly, we found that AlphaFold reliably predicted the presence of a second closure motif, which we term CM2, within SYCP2 at residues 466-484 (Figure 3B and C). Based on our previous work, we are very aware that AlphaFold will frequently insert unstructured regions of polypeptide into a HORMA domain as a “pseudo-CM” (Kane et al., 2025). We thus took multiple approaches to confirm the validity and functionality of SYCP2-CM2. Firstly, multiple sequence alignment of SYCP2 sequences from various vertebrates shows the high degree of conservation of the novel interaction motif (Supplementary Figure 4). Critically, CM2 contains the sequence KKMSE that is also described for closure motifs of HORMAD1 and 2 (Figure 3D), making CM2 more similar to the HORMAD *cis-*closure motifs.

**Figure 3.**
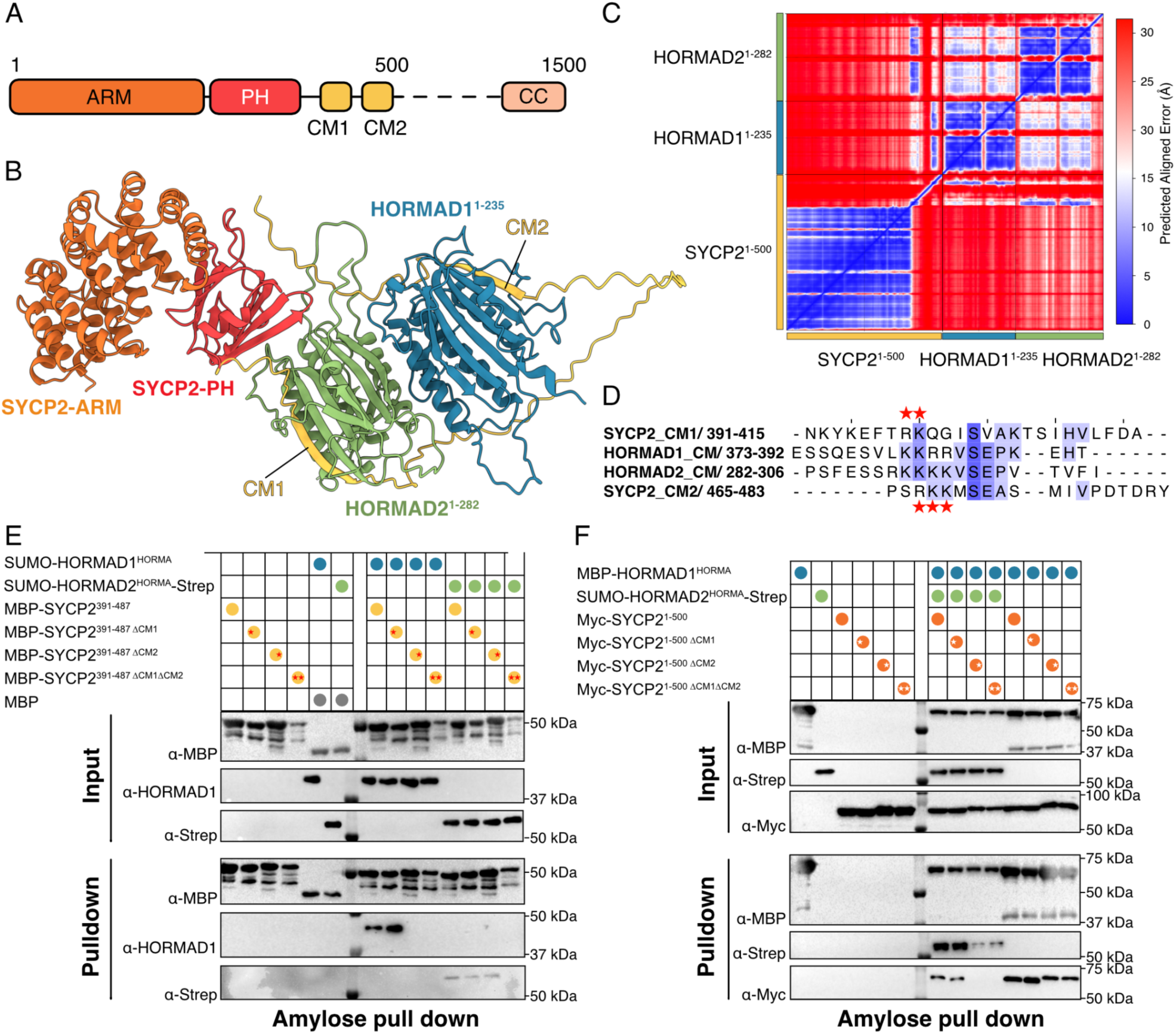
HORMAD1 and HORMAD2 bind via distinct closure motifs in SYCP2. A) Domain schematic of mouse SYCP2. ARM = ARM-like domain; PH = pleckstrin homology domain (also referred to as SLD; Spt16M-like domain); CM1 = closure motif 1; CM2 = closure motif 2; CC = coiled-coil. The dotted line refers to residues 501-1325, which are predicted to be unstructured and are not shown here. B) Cartoon representation of the AlphaFold2 prediction of the SYCP2^1-500^ / HORMAD1^HORMA^ / HORMAD2^HORMA^ complex. SYCP2 is coloured as in A). C) Predicted alignment error (PAE) plot of the prediction shown in B). D) Comparison of the closure motifs for SYCP2 CM1 (top), HORMAD1 CM, HORMAD2 CM, and SYCP2 CM2. E) Amylose beads pull down on MBP-SYCP2 closure motif constructs against HORMAD1^HORMA^ or HORMAD2^HORMA^ as indicated. ΔCM1 indicates the R398A K399A mutation, ΔCM2, the R468A K469A K470A mutation. Anti-HORMAD1 was used to detect HORMAD1 in the input and the pull down fractions; anti-Strep was used to detect Strep-tagged HORMAD2^HORMA^. F) Amylose pull down on MBP-HORMAD1^HORMA^ against Myc-SYCP2^1-500^ constructs coexpressed in the absence or presence of HORMAD2. The ΔCM mutations used are the same as in E).

To test the role of CM2 we performed a pull down experiment from bacteria coexpressing HORMAD1^HORMA^, HORMAD2^HORMA^ and residues 391-487 of SYCP2 with an N-terminal MBP fusion (MBP-SYCP2^391-487^). This region of SYCP2 should contain both closure motifs, the previously described CM1 and the hypothetical CM2. We created constructs where these CMs were mutated at the conserved R/K residues to alanine, so ΔCM1 = R398A/K399A and ΔCM2 = R468A/K469A/K470A. We prepared different versions of MBP-SYCP2^391-487^, 6xHis-SUMO-HORMAD1^HORMA^, 6xHis-SUMO-HORMAD2^HORMA^-Strep (Supplementary Figure 1). In the pull down assay the levels of MBP-SYCP2 pulled down on the amylose beads were consistent for the different MBP-SYCP2^391-487^ constructs (Figure 3E, pull down). We found that the wildtype MBP-SYCP2 could pull down both HORMAD1^HORMA^ and HORMAD2^HORMA^, confirming the presence of an interaction motif for both HORMADs within this region of SYCP2. MBP-SYCP2^391-487ΔCM1^ still interacted with HORMAD1^HORMA^, but the mutation of CM2 specifically disrupted this interaction, confirming the role of CM2, and and showing HORMAD1 specificity for CM2 in the assayed condition and SYCP2 fragment. Curiously, single mutations of CM1 and CM2 showed the same levels of interaction for HORMAD2^HORMA^, suggesting that HORMAD2, in addition to binding to CM1, is also able to bind to CM2. The double mutation - MBP-SYCP2^391-487ΔCM1ΔCM2^ resulted in a loss of detectable binding for both HORMAD1^HORMA^ and HORMAD2^HORMA^ (Figure 3E).

We next asked what role the ARM and PH domains of SYCP2 might play in assembling a complex, and whether HORMAD1 and HORMAD2 can bind simultaneously to SYCP2. We carried out a similar bacterial coexpression experiment as described above, but in this case the bait was MBP-HORMAD1^HORMA^, and we used an N-terminally Myc-tagged version of SYCP2 encompassing residues 1-500 (Myc-SYCP2^1-500^) (Figure 3F). The mutations in the closure motifs are as described above. We found that both HORMAD2 and SYCP2^1-500^ bind to MBP-HORMAD1 when both closure motifs are present. A mutation in the HORMAD2 binding CM1 does not reduce the level of HORMAD2 pulled down. When CM2 is mutated (either alone or together with CM1), we could no longer detect SYCP2 in the pull down, but did observe some HORMAD2, presumably via interaction with HORMAD1 (Figure 3F). In the absence of HORMAD2, we observed a different behaviour for HORMAD1. Here we observed, as expected, a robust interaction between HORMAD1 and SYCP2 in the presence of a functional CM2. However, once CM2 was mutated, there was still a substantial band corresponding to Myc-SYCP2^1-500^, even when both CM1 and CM2 were mutated (Figure 3F). We therefore conclude that in the absence of HORMAD2, HORMAD1 can bind to the ARM-PH domain of SYCP2.

### Characterisation of the SYCP2-HORMAD1-HORMAD2 complex

Having shown that HORMAD1, HORMAD2 and SYCP2 can all bind together, we sought to gain further insights into the complex. We coexpressed and purified the complex 6xHis-Myc-SYCP2^1-500^/6xHis-MBP-HORMAD1^HORMA^/6xHis-SUMO-HORMAD2^HORMA^-Strep for further analysis, using MBP-trap affinity chromatography followed by anion-exchange chromatography and size-exclusion chromatography (Figure 4A). Employing mass photometry, we observed three distinct peaks at 118, 170 and 233 kDa. 118 kDa either corresponds to a heterodimer of two HORMAD1^HORMA^-HORMAD2^HORMA^ (theoretical MW of 119 kDa) or a heterodimeric complex of HORMAD2^HORMA^-SYCP2^1-500^ (theoretical MW of 118 kDa). The 170 kDa peak likely corresponds to a hetero-trimeric complex of SYCP2^1-500^-HORMAD1^HORMA^-HORMAD2^HORMA^ (theoretical mass of 187 kDa). We propose that the 233 kDa peak is 1 molecule of SYCP2^1-500^ and 3 molecules of HORMA domain. We suggest that the complex contains 2xHORMAD2^HORMA^, where one binds via the closure motif of SYCP2 and another through the interaction interface with HORMAD1^HORMA^.

**Figure 4.**
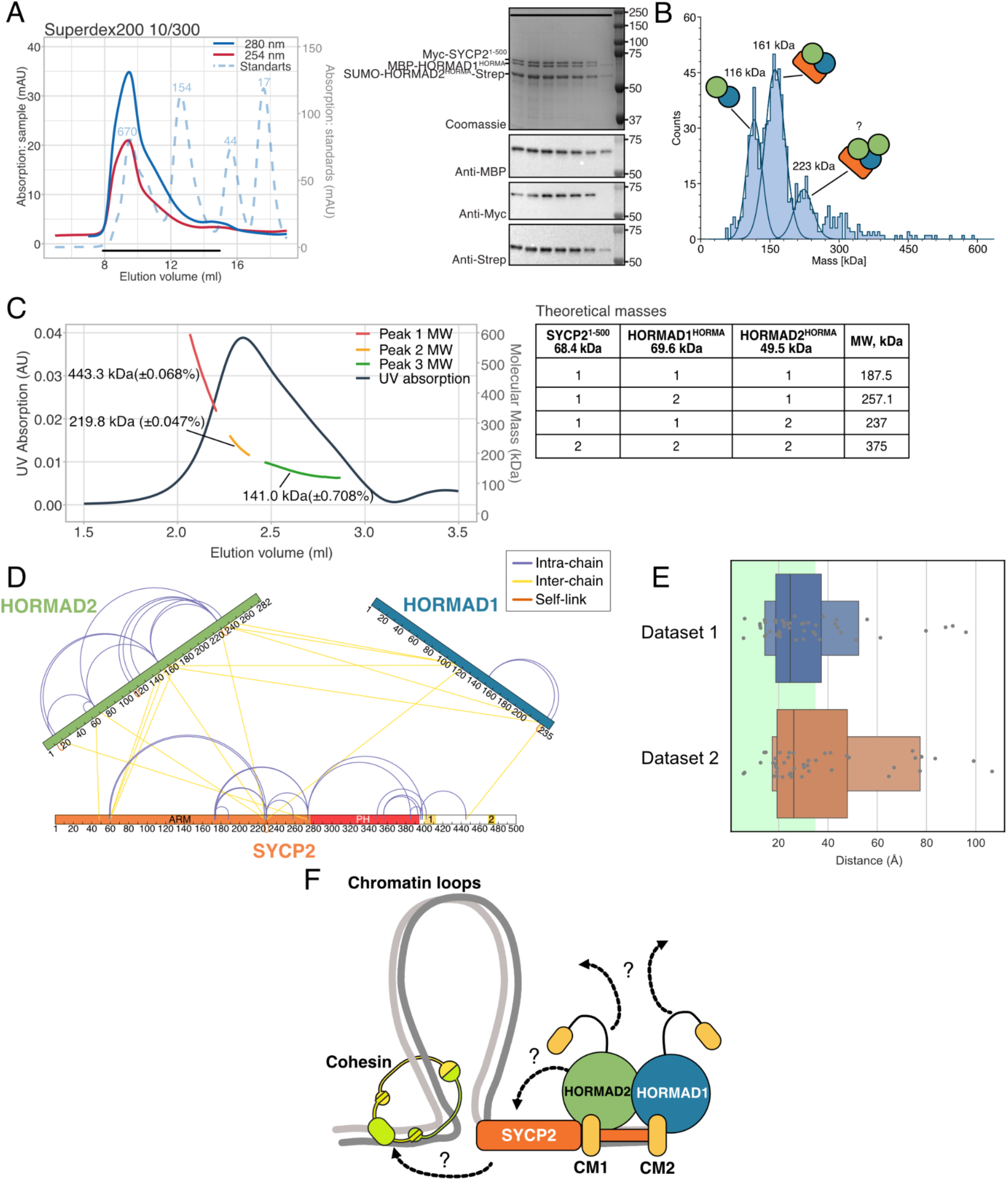
Characterisation of the SYCP2-HORMAD1-HORMAD2 complex. A) SEC profile of the purification of Myc-SYCP2^1-500^/ MBP-HORMAD1^HORMA^/ SUMO-HORMAD2^HORMA^-Strep. The samples on the SDS-PAGE gel correspond to the black bar indicated in the chromatogram. B) Mass photometry data recorded for the purified Myc-SYCP2^1-500^/ MBP-HORMAD1^HORMA^/ SUMO-HORMAD2^HORMA^-Strep complex after size-exclusion chromatography. Samples were recorded for 60 secs at ∼30 nM concentration. C) SEC-MALS analysis of the Myc-SYCP2^1-500^/ MBP-HORMAD1^HORMA^/ SUMO-HORMAD2^HORMA^-Strep complex. Samples were measured at ∼10 μM concentration on the column Superdex200 5/150 (Cytiva) in 20 mM HEPES pH 8.2, 300 mM NaCl, 1 mM TCEP. D) XL-MS dataset on the purified Myc-SYCP2^1-500^/ MBP-HORMAD1^HORMA^/ SUMO-HORMAD2^HORMA^-Strep complex. Samples were cross-linked with DSBU, trypsinated, and analysed. Data visualised in XiVis. Cross-links coloured as indicated in the legend. E) Distance distribution of cross-links from two independent datasets, modelled onto the top-scoring SYCP2^1-500^/HORMAD1^HORMA^/HORMAD2^HORMA^ predicted structure (from Figure 3B and C). The green box corresponds to the Cα-Cα cut-off distance of 35Å. F) SYCP2 contains two closure motifs, with CM2 preferentially bound by HORMAD1. HORMAD1 and HORMAD2 form a heterodimer via their HORMA domains, which potentially leaves their closure motifs (yellow) exposed and able to bind to further partners; either more HORMA domains or additional factors.

Given that mass photometry is carried out at low concentrations, we followed up our analyses with size-exclusion chromatography coupled with multi-angle light scattering (SEC-MALS) (Figure 4C). Similar to the SEC-MALS on HORMAD1^HORMA^ and HORMAD2^HORMA^, we observed a contiguous peak from which we derived three mass measurements; 141 kDa (green trace), 219.8 kDa (yellow trace) and 444.3 kDa (red trace). With a complex of 1:1:1 stoichiometry having a theoretical molecular mass of 187.5 kDa, we assume we have a mixture of species, including higher-order complexes. What components of the SYCP2-HORMAD1^HORMA^-HORMAD2^HORMA^ complex might be driving the formation of higher-order assemblies? Above, we showed that the HORMA domains will form dimers, likewise, the SYCP2 ARM-PH region is well characterised and appears to intrinsically form monomers (Feng et al., 2017). Our focus therefore, landed on the relationship between SYCP2 and the HORMADs. We purified the complex, containing the closure motifs region of SYCP2 (SYCP2^391-487^) with N-terminal Myc-fusion, HORMAD1^HORMA^ with N-terminal SUMO tag and C-terminal StrepII-tag and HORMAD2 with N-terminal SUMO tag (Supplementary Figure 5A). We measured the mass of this complex with SEC-MALS (Supplementary Figure 5B). Consistent with the complex containing SYCP2^1-500^ we observed masses corresponding to higher order complexes, suggesting that the relationship between the HORMA domains and the SYCP2 CMs drive oligomerisation. We explored whether AlphaFold2 or AlphaFold3 might be able to produce a high-confidence prediction with the variations of SYCP2 HORMAD1^HORMA^ and HORMAD2^HORMA^. However, the only structure that was predicted with confidence was a 1:1:1 stoichiometry, as shown in Figure 3C.

Finally, we analysed the structural organisation of the complex using crosslinking disuccinimidyl dibutyric urea (DSBU) coupled with mass-spectrometry (Figure 4D). We observed multiple cross-links between HORMAD1 and HORMAD2, and between HORMAD2 and the ARM domain of SYCP2. The only cross-link proximal to the closure motif region of SYCP2 and a HORMAD showed a link between HORMAD1 and the region between CM1 and CM2. How well does the XL-MS data support the AlphaFold model of the SYCP2^1-500^-HORMAD1^HORMA^-HORMAD2^HORMA^ complex? We modelled the experimental crosslinks onto the AlphaFold prediction. We found that the majority of the cross-links corresponded well to the predicted complex, across two independent datasets, due to their distances below the maximum allowed 35Å Cα-Cα distance (Figure 4E, green box). For those cross-links that are not consistent with the model, we suspect that this is either due to the higher stoichiometry discussed above, or additional flexibility/contacts within the complex, for example, an interaction between HORMAD1 and SYCP2.

## Discussion

Our study set out to address the question: what is the interplay between SYCP2, HORMAD1 and HORMAD2? Previous work has shown that HORMAD1 and HORMAD2 interact with one another, with the C-terminal closure motif of HORMAD2 shown to interact with HORMAD1^HORMA^ via pull down experiments (West et al., 2019) and through co-crystallisation (Wang et al., 2023). Recent yeast two-hybrid data further suggested that the HORMA domains of HORMAD1 and HORMAD2 could also interact (Raveendran et al., 2026). We followed this up and conclusively showed, through biophysical and hybrid structural biological analysis, that the HORMA domains of HORMAD1 and HORMAD2 form heterodimers, independently of closure motifs. While many HORMAs form both homo- and heterodimers via their HORMA domain, meiotic HORMAs have only previously been shown to interact via closure motifs, and not via their HORMA domains (Kim et al., 2014; Wang et al., 2023; West et al., 2017). We identify a conserved interface that favours heterodimer formation, but does not exclude homodimer formation. Several aspects remain unclear; what is the affinity of the interface? What is the topology of the HORMA domains in the heterodimer? How does TRIP13 interact with a heterodimer of HORMAD1-HORMAD2?

SYCP2 has been shown to directly recruit HORMAD2 via a closure motif in the region of 391-415, which we term here CM1 (West et al., 2019). AlphaFold modelling suggested that a second closure motif might also exist in SYCP2 in residues 466-484, which preferentially binds to HORMAD1. Through pull downs and mutational analysis, we find disruptions to CM2 specifically ablate the ability of SYCP2^391-487^ to bind to HORMAD1. Curiously, we find that HORMAD2 also has an ability to bind to CM2, since HORMAD2 binding is only lost when both SYCP2 closure motifs are mutated. A caveat is that the assayed CM mutation may only partially disable HORMAD binding when they are introduced in isolation. Yet, the combination of these mutations may cause misfolding of the SYCP2 fragment, which may be reflected by low protein stability, and may lead to a non-specific loss of HORMAD binding. These features of the experimental system may explain the apparent divergence between in vivo and in vitro requirements for HORMAD2-SYCP2 interactions. This suggests that HORMAD2 might be a promiscuous CM binder, whereas HORMAD1 may have clear sequence preferences. When using a longer region of SYCP2, containing the ARM-PH domains, we found that HORMAD1 will bind to this construct, even when both SYCP2 closure motifs are mutated. We presume that this is down to an interaction between the HORMA domain of HORMAD1 and the ARM-PH domain of SYCP2. We have recently shown an equivalent interaction for the HORMA domain of Hop1 and the ARM-PH domain of Red1 in budding yeast (Chen et al., 2025). When HORMAD2^HORMA^ is included in the experiment, we no longer see an interaction between HORMAD1^HORMA^ and SYCP2^1-500^ in the absence of CM2. Therefore, the ability of HORMAD1 to interact with the ARM-PH domain of SYCP2 depends on HORMAD2. We speculate that this represents different states of assembly, and that one function of the ARM-PH region of SYCP2 might be in facilitating the proper organisation of the HORMAD1 and HORMAD2 on the axis.

The previous model for meiotic axis assembly postulates that HORMAD2 is recruited to SYCP2 via the SYCP2 closure motif. In binding HORMAD2 would then release its own closure motif that could be bound by HORMAD1. Our work here proposes some modifications to that model. Firstly, we find that HORMAD1 will bind to a second closure motif (CM2) in SYCP2. Under specific conditions SYCP2 CM2 may also bind to HORMAD2, but we speculate that the ARM-PH domains of SYCP2 might promote a particular organisation of the HORMADs, whereby HORMAD2 and HORMAD1 engage CM1 and CM2 on the axis, respectively (Figure 4F). Secondly, we find that HORMAD1 and HORMAD2 form a heterodimer between the HORMA domains, independently of their own closure motifs (Figure 1C-E, Figure 2). Furthermore, in the presence of full-length proteins, two further CMs become available (the *cis*-CMs of HORMAD1 and HORMAD2) several distinct types of HORMAD assemblies may form on meiotic axes. Indeed, while we did not see any direct evidence of homodimerisation of the HORMADs, the side-chain chemistry of the HORMA domains would not preclude this from happening. This adds another level of potential diversity to the nature of the assembly of the meiotic axis, behavior that might be connected to the diverse roles of HORMAD1 and HORMAD2 in meiotic DSB formation, DNA damage signalling, and the subsequent controlled removal of these factors during synapsis.

## Methods

### Yeast two-hybrid analysis

Proteins were cloned into pGAD and pGBDU vectors for the yeast two-hybrid assay. The obtained plasmids were co-transformed into *S. cerevisiae* reporter strain yWL365 (kindly provided by Gerben Vader), and the cells were plated onto selective mediums without leucine and uracil (-LU). For the drop assay, transformed cells with OD600 0.5 were dropped with serial dilutions of 1:10, 1:100 and 1:1000 on the plates with selective medium lacking leucine and uracil (-LU) as a control, and without leucine, uracil and histidine (-LUH) to analyze the interactions. -LUH plates were incubated for 4 days at 30°C and -LU plates at 25°C for 2 days.

### Protein Purification

All vectors used here are part of our InteBac Vector Suite (Altmannova et al., 2021). The plasmids used in this study are listed in the Supplementary Table 1. Genes for mouse HORMAD1, HORMAD2 and SYCP2 were used from synthetic geneblocks (IDT), without codon optimisation.

Both *<u>HORMAD1^HORMA^</u> and <u>HORMAD2^HORMA^</u>* with 3C HRV cleavable N-terminal tags: for HORMAD1 - 6xHis-MBP tag, for HORMAD2 - N-terminal 6xHis-SUMO and C-terminal STREPII-tag were expressed in chemically competent E.coli BL21(DE3)Star cells. Protein expression was induced by adding 250 µM IPTG at 16°C overnight. Cell pellets were resuspended in buffer A (50 mM HEPES pH 8.2, 300 mM NaCl, 5% Glycerol, 1 mM MgCl_2_, 0.01% TritonX100, 1 mM βME, 5 mM imidazole, benzonase and AEBSF protease inhibitor) and lysed by sonication. Cell lysates were centrifuged at 10 000 rpm, 4°C for 1 hour. Cleared lysate was loaded on a 5 ml affinity column (MBP-trap (Cytiva) for HORMAD1^HORMA^ and Streptactin®XT column (IBA) for HORMAD2^HORMA^). After loading, the columns were washed with 10 column volumes of Buffer A followed by 3 column volumes of Buffer A + 1 mM ATP. HORMAD1^HORMA^ was eluted by 10 column volumes of Buffer A + 50 mM maltose. HORMAD2^HORMA^ was eluted by 10 column volumes of Buffer A + 50 mM biotin. The protein fractions with the highest absorption at 280 nm were combined and loaded onto the ResourceQ anion-exchange column (Cytiva), preequilibrated with Buffer B (50 mM HEPES pH 8.2, 150 mM NaCl, 5% Glycerol, 1 mM TCEP). After the loading of the sample, the column was washed with 5 column volumes of buffer B, followed by gradient elution increasing the concentration of NaCl up to 1 M. The fractions containing HORMAD1^HORMA^ or HORMAD2^HORMA^ were loaded onto Superdex200 16/600 and Superdex200 10/300 (Cytiva), respectively, preequilibrated with 50 mM HEPES pH 8.2, 300 mM NaCl, 5% Glycerol, 1 mM TCEP. The fractions containing HORMAD1^HORMA^ or HORMAD2^HORMA^ were concentrated on Pierce protein concentrators (30K MWCO, ThermoFisher), aliquoted and stored at -70°C.

Separately, the *<u>HORMAD1^HORMA^ with N-terminal 6xHis-SUMO</u>* fusion was expressed and purified using the same protocol, with an exception at the affinity chromatography step. There, the cleared lysate was loaded onto a 5 ml Talon column (Cytiva), followed by a wash with 10 column volumes of Buffer B + 20 mM imidazole and 3 column volumes of Buffer B + 1 mM ATP. The protein was eluted with imidazole gradient up to 250 mM. The subsequent purification steps were the same as for the purifications of individual HORMAD1^HORMA^ and HORMAD2^HORMA^.

*<u>SYCP2^391-487^ and corresponding CM mutants (CM1, CM2, CM1CM2)</u>* were produced with cleavable 3C HRV N-terminal MBP tag in chemically competent *E. coli* BL21(DE3)Star cells. Protein expression was induced by adding 250 µM IPTG at 16°C overnight. Cell pellets were resuspended in the buffer C (50 mM HEPES pH 7.5, 300 mM NaCl, 5% Glycerol, 1 mM MgCl_2_, 1 mM βME, benzonase and AEBSF protease inhibitor) and lysed by sonication. Cell lysates were centrifuged at 4°C and 10000 rpm for 1 h. Cleared lysates were loaded onto 5 ml MBP-trap column (Cytiva), followed by a wash with 10 column volumes of buffer C, 3 column volumes of Buffer C + 1 mM ATP and eluted with 10 column volumes of buffer C + 50 mM maltose. Fractions containing *SYCP2^391-487^* were loaded on Superdex200 16/600 column (Cytiva), and after the chromatography, they were concentrated on Pierce protein concentrators (10K MWCO, ThermoFisher), aliquoted and stored at -70°C.

*<u>SYCP2^1-500^</u>* was produced with a cleavable 3C HRV N-terminal Myc tag in chemically competent *E. coli* BL21(DE3)Star cells. Protein expression was induced by adding 250 µM IPTG at 16°C overnight. Cell pellets were resuspended in the buffer E (50 mM HEPES pH 7.5, 300 mM NaCl, 5% Glycerol, 1 mM MgCl_2_, 1 mM βME, 5 mM imidazole, benzonase and AEBSF protease inhibitor) and lysed by sonication. Cell lysates were centrifuged at 4°C and 10000 rpm for 1 h. Cleared lysates were loaded onto 5 ml His-trap column (Cytiva), followed by a wash with 10 column volumes of buffer E + 20 mM imidazole, 3 column volume of Buffer E + 20 mM imidazole + 1 mM ATP and eluted with gradient of imidazole up to 250 mM. Fractions containing *SYCP2^1-500^* were loaded on Superdex200 10/300 column (Cytiva) and after the chromatography they were concentrated on Pierce protein concentrators (30K MWCO, ThermoFisher), aliquoted and stored at -70°C.

#### Coexpression of *HORMAD1^HORMA^ and HORMAD2^HORMA^*

The plasmids with HORMAD1^HORMA^ with N-terminal MBP-tag and HORMAD2^HORMA^ with N-terminal 6xHis-SUMO and C-terminal StrepII tags were co-transformed into chemically competent *E.coli* BL21(DE3)Star cells. The protein expression and purification protocol was identical to the protocol for individual HORMAD1^HORMA^ and HORMAD2^HORMA^.

#### Coexpression of SYCP2^1-500^, *HORMAD1^HORMA^ and HORMAD2^HORMA^*

The plasmids used for the expression of 6xHis-MBP-HORMAD1^HORMA^, 6xHis-SUMO-HORMAD2^HORMA^-Strep and 6His-Myc-SYCP2^1-500^ were cotransformed into chemically competent *E.coli* BL21(DE3)Star cells. Protein expression was induced by adding 250 µM IPTG at 16°C overnight. Cell pellets were resuspended in the buffer F (50 mM HEPES pH 7.5, 300 mM NaCl, 5% Glycerol, 1 mM MgCl_2_, benzonase and AEBSF protease inhibitor) and lysed by sonication. Cell lysates were centrifuged at 4°C and 10000 rpm for 1 h. Cleared lysates were loaded onto 1 ml Strep-trap (Cytiva). After loading, the column were washed with 10 column volumes of Buffer F followed by 3 column volumes of Buffer F + 1 mM ATP, and eluted by 10 column volumes of Buffer F + 50 mM biotin. The fractions with the highest absorption at 280 nm were combined and loaded onto the ResourceQ anion-exchange column (Cytiva), preequilibrated with Buffer H (50 mM HEPES pH 7.5, 100 mM NaCl, 5% Glycerol, 1 mM TCEP). After the loading of the sample, the column was washed with 5 column volumes of buffer H, followed by gradient elution increasing the concentration of NaCl up to 1 M. The fractions containing SYCP2^1-500^, HORMAD1^HORMA^ and HORMAD2^HORMA^ were loaded onto Superdex200 10/300 (Cytiva) preequilibrated with 50 mM HEPES pH 7.5, 300 mM NaCl, 5% Glycerol, 1 mM TCEP. Fractions containing SYCP2^1-500^, HORMAD1^HORMA^ and HORMAD2^HORMA^ were loaded on Superdex200 10/300 column (Cytiva) and after the chromatography they were concentrated on Pierce protein concentrators (30K MWCO, ThermoFisher), aliquoted and stored at -70°C.

#### Coexpression of SYCP2^391-487^, *HORMAD1^HORMA^ and HORMAD2^HORMA^*

The plasmids with 6xHis-SYCP2^391-487^, 6xHis-SUMO-HORMAD1^HORMA^-Strep and 6His-SUMO-HORMAD2^HORMA^ were cotransformed into chemically competent *E.coli* BL21(DE3)Star cells. The protein expression and purification protocol was identical to the protocol of coexpression of SYCP2^1-500^, HORMAD1^HORMA^ and HORMAD2^HORMA^ with an exception on the size-exclusion chromatography step where the Superdex 200 16/600 (Cytiva) was used.

### Coexpression IP

Proteins for coexpression IP were produced in chemically competent E. coli BL21(DE3)Star cells. Protein expression was induced by 500 µM IPTG at 37°C for 3 hours. Cell pellets were resuspended in 2 ml of pull-down buffer (20 mM HEPES pH 7.5, 300 mM NaCl, 5% Glycerol, 1 mM MgCl_2_, 1 mM βME, 5 mM imidazole) and lysed by sonication. Lysates were centrifuged at 10000 rpm for 1 hour at 4°C. Cleared lysates were incubated with preequilibrated in pull down buffer and preblocked with 5 mg/ml BSA Amylose beads (10 μl per reaction) for 30 min with rotation at 8°C. The beads were washed with 200 μl of pull down buffer three times and the proteins were eluted with 40 μl of elution buffer with 50 mM maltose. The samples were mixed with 6x Laemmli buffer, boiled at 95°C for 3 min and loaded on 10% SDS-PAGE. After the electrophoresis gels were stained with Der Blaue Jonas gel dye (GRP) or used for the western blotting and immunostaining.

### Mass Photometry

The proteins for mass photometry analysis were diluted in degased and filtered through 0.2 μm filter MP buffer (20 mM HEPES pH 7.5, 100 mM NaCl, 5% Glycerol, 1 mM TCEP) immediately before the measurement. The analysis of the measurements were performed in DiscoverMP software using MP1 standards (Refeyn), diluted in the same buffer, for the calibration. All measurements were performed on OneMP instrument (Refeyn).

### AlphaFold predictions

AlphaFold 2.3.1 (Evans et al., 2021; Jumper et al., 2021) and AlphaFold 3.0.1 (Abramson et al., 2024) were installed on the Raven compute cluster at the Max Planck Computing and Data Facility (MPCDF). For all analyses described, 25 runs were carried out in total, based on 5 initial seeds. Multiple sequence alignments were automatically generated using the JackHMMER (Johnson et al., 2010). Custom scripts were used to generate PAE plots and to compare the resulting ipTM scores between runs.

### Cross-linking Mass Spectrometry (XL-MS)

Proteins at concentration of 4 μM in the buffers after size-exclusion chromatography were mixed with 4 μl of DSBU (200 mM) and incubated for 1 hour at 25°C. The reaction was stopped by adding 20 μL of 1 M Tris pH 8.5 and incubated for 30 min in 25°C. The sample was mixed with 4 times the volume of cold 100% acetone and incubated overnight at -20°C. The following analysis was performed as described in (Pan et al., 2018). Cross-links were analysed and visualised in XiView (Combe et al., 2024).

### Pull downs

For the pull down with SYCP2^391-487^ wild type and CM mutants (9 μM) and HORMAD1^HORMA^ / HORMAD2^HORMA^ (6 μM), the proteins were mixed in the reaction buffer (20 mM HEPES pH 7.5, 150 mM NaCl, 0.025% Tween20, 10% Glycerol, 1 mM TCEP) and incubated on ice for 1 hour. Pre-equilibrated in the reaction buffer and preblocked with 5 mg/ml BSA amylose beads were added to the reactions (10 μl per reaction) and incubated for 30 min in the thermomixer at 8°C and 950 rpm. The beads were then washed with 200 μl of reaction buffer three times and the proteins were eluted by adding 40 μl of reaction buffer + 50 mM maltose. The samples were mixed with 6x Laemmli buffer, boiled at 95°C for 3 min and loaded on 10% SDS-PAGE. After the electrophoresis gels were stained with Der Blaue Jonas gel dye or used for the western blotting and immunostaining.

For the pull down with HORMAD1^HORMA^ (3 μM) and HORMAD2^HORMA^ (6 μM), the proteins were mixed and incubated in the reaction buffer at 8°C for 24 hours. Pre-equilibrated in the reaction buffer and preblocked with 5 mg/ml BSA Streptactin beads were added to the reactions (10 μl per reaction) and incubated for 30 min in the thermomixer at 8°C and 950 rpm. The beads were then washed with 200 μl of reaction buffer three times and the proteins were eluted by adding 40 μl of reaction buffer + 50 mM biotin. The samples were mixed with 6x Laemmli buffer, boiled at 95°C for 3 min and loaded on 10% SDS-PAGE. After the electrophoresis gels were stained with Der Blaue Jonas gel dye or used for the western blotting and immunostaining.

### Western Blotting

For the immunostaining analysis, the samples were run on 10% SDS-PAGE gels and transferred onto a nitrocellulose membrane with semi-dry western blotting. After the transfer, the membranes were blocked with 5% nonfat milk in PBST (1xPBS buffer with 0.1% Tween20) and then incubated with specific antibodies in the dilution 1:1000. The following antibodies were used: for HORMAD1^HORMA^ the anti-HORMAD1 (rabbit, Abcam, #ab178432), for HORMAD1^HORMA^ and for SYCP2^391-487^ wild type and CM mutants anti-MBP (rabbit, NEB, #E8032L), for HORMAD2^HORMA^ anti-Strep (mouse, Cube-Biotech, #40070), for SYCP2^1-500^ anti-Myc (mouse, BioLegend, #658502). After the incubation with primary antibodies, the membranes were washed with PBST and incubated with the secondary goat anti-rabbit (Merck, 401353) or goat anti-mouse (Merck, 401215) IgG peroxidase conjugate antibody. The proteins were visualized by chemiluminescence detection (WesternBright ECL, Advansta).

### SEC-MALS

50 μl of sample containing ∼10 μM of protein was run on the column Superdex 200 5/150 (for HORMAD1^HORMA^-HORMAD2^HORMA^ heterodimer and SYCP2^391-487^-HORMAD1^HORMA^- HORMAD2^HORMA^ complex) in 20 mM HEPES pH 8.2, 150 mM NaCl, 1 mM TCEP or Superose6 5/150 (for the SYCP2^1-500^-HORMAD1^HORMA^- HORMAD2^HORMA^ complex) in 20 mM HEPES pH 8.2, 300 mM NaCl, 1 mM TCEP with flowrate 0.3 ml/min on 1260 Infinity II LC System (Agilent). MALS was measured using a Wyatt DAWN detector combined with the size exclusion column.

## Acknowledgements

The authors would like to thank Rahmiye Kürkcü, Jennifer Jüngling and Susanne Astrinidis for their help with the cloning of the constructs used in this study; Gerben Vader, Atilla Toth and Veronika Altmannova for the proofreading the manuscript, insightful discussions and constructive feedback; Veronika Altmannova for the help with establishing yeast two-hybrid assays. This work was funded by Max Planck Society. ES thanks the International Max Planck Research School ‘From Molecules to Organisms’.

