## Supplementary Data for "Assembly principles of a SYCP2-HORMAD1-HORMAD2 mammalian meiotic axis complex"

Supplementary Table 1 - Plasmids used in this study

| Plasmid ID | Purpose | Backbone | Resistance and Origin | Description | Reference |
|---|---|---|---|---|---|
| pWL2349 | Y2H | pGAD-C1 | Amp/Ori | AD-HORMAD1$^{1-235}$ | This study |
| pWL2353 | Y2H | pGBDU-C1 | Amp/Ori | BD-HORMAD1$^{1-235}$ | This study |
| pWL2350 | Y2H | pGAD-C1 | Amp/Ori | AD-HORMAD2$^{1-282}$ | This study |
| pWL2354 | Y2H | pGBDU-C1 | Amp/Ori | BD-HORMAD2$^{1-282}$ | This study |
| pWL1565 | Y2H | pGAD-C1 | Amp/Ori | Empty vector | Altmannova, Firlej et al., NAR, 2023 |
| pWL1564 | Y2H | pGBDU-C1 | Amp/Ori | Empty vector | Altmannova, Firlej et al., NAR, 2023 |
| pWL2484 | Bacterial Expression | pRSF | Kan/Ori | 6xHis-SUMO-HORMAD2$^{1-282}$ | This study |
| pWL2230 | Bacterial Expressio | pCDF | Spec/Ori | 6xHis-Myc-SYCP2$^{391-487}$ | This study |

| | | | | | |
|---|---|---|---|---|---|
| | n | | | | |
| pWL2299 | Bacterial Expression | pCOLI | Amp/Ori | 6xHis-SUMO-HORMAD1$^{1-235}$-StrepII | This study |
| pWL2643 | Bacterial Expression | pRSF | Kan/Ori | 6xHis-MBP-HORMAD1$^{1-235}$ | This study |
| pWL1397 | Bacterial Expression | pCOLI | Amp/Ori | 6xHis-SUMO-HORMAD1$^{1-235}$ | Dereli et al. Nat. Com., 2024 |
| pWL1699 | Bacterial Expression | pCOLI | Amp/Ori | 6xHis-SUMO-HORMAD2$^{1-282}$-StrepII | This study |
| pWL2300 | Bacterial Expression | pCOLI | Amp/Ori | 6xHis-MBP-SYCP2$^{391-487}$ | This study |
| pWL2307 | Bacterial Expression | pCOLI | Amp/Ori | 6xHis-MBP-SYCP2$^{391-487CM1}$ | This study |
| pWL2308 | Bacterial Expression | pCOLI | Amp/Ori | 6xHis-MBP-SYCP2$^{391-487CM2}$ | This study |
| pWL2312 | Bacterial Expressio | pCOLI | Amp/Ori | 6xHis-MBP-SYCP2$^{391-487CM1CM2}$ | This study |

| | | | | | |
|---|---|---|---|---|---|
| | n | | | | |
| pWL2509 | Bacterial Expression | pCDF | Spec/Ori | 6xHis-Myc-SYCP2$^{1\text{-}500}$ | This study |
| pWL2773 | Bacterial Expression | pCDF | Spec/Ori | 6xHis-Myc-SYCP2$^{1\text{-}500\ CM1}$ | This study |
| pWL2774 | Bacterial Expression | pCDF | Spec/Ori | 6xHis-Myc-SYCP2$^{1\text{-}500\ CM2}$ | This study |
| pWL2775 | Bacterial Expression | pCDF | Spec/Ori | 6xHis-Myc-SYCP2$^{1\text{-}500\ CM1CM2}$ | This study |

Supplementary Table 2 - Yeast strain used in this study

| | | |
|---|---|---|
| yWL365 | MATa, ura3-52, leu2-3, his3, trp1, gal4del, gal80del,<br>GAL2-ADE2, LYS2::GAL1-HIS3, met2::GAL7-lacZ | Gerben Vader |

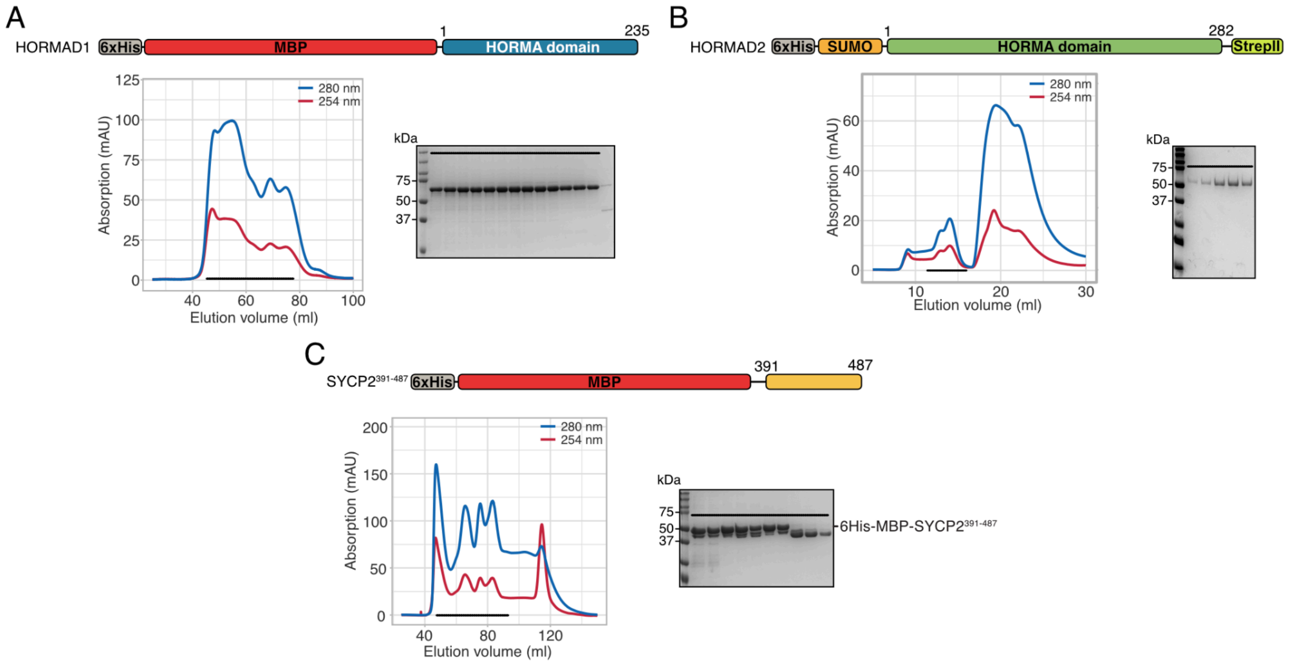


*Supplementary Figure 1 - Representative Protein Purifications*

Upper panels: cartoon representations of protein fragment purified; lower panels: left - size-exclusion chromatography profiles, right - SDS-PAGE of the fractions containing the purified proteins. A: HORMAD1$^{HORMA}$ purification, B: HORMAD2$^{HORMA}$ purification, C: MBP-SYCP2$^{391\text{-}487}$ purification.

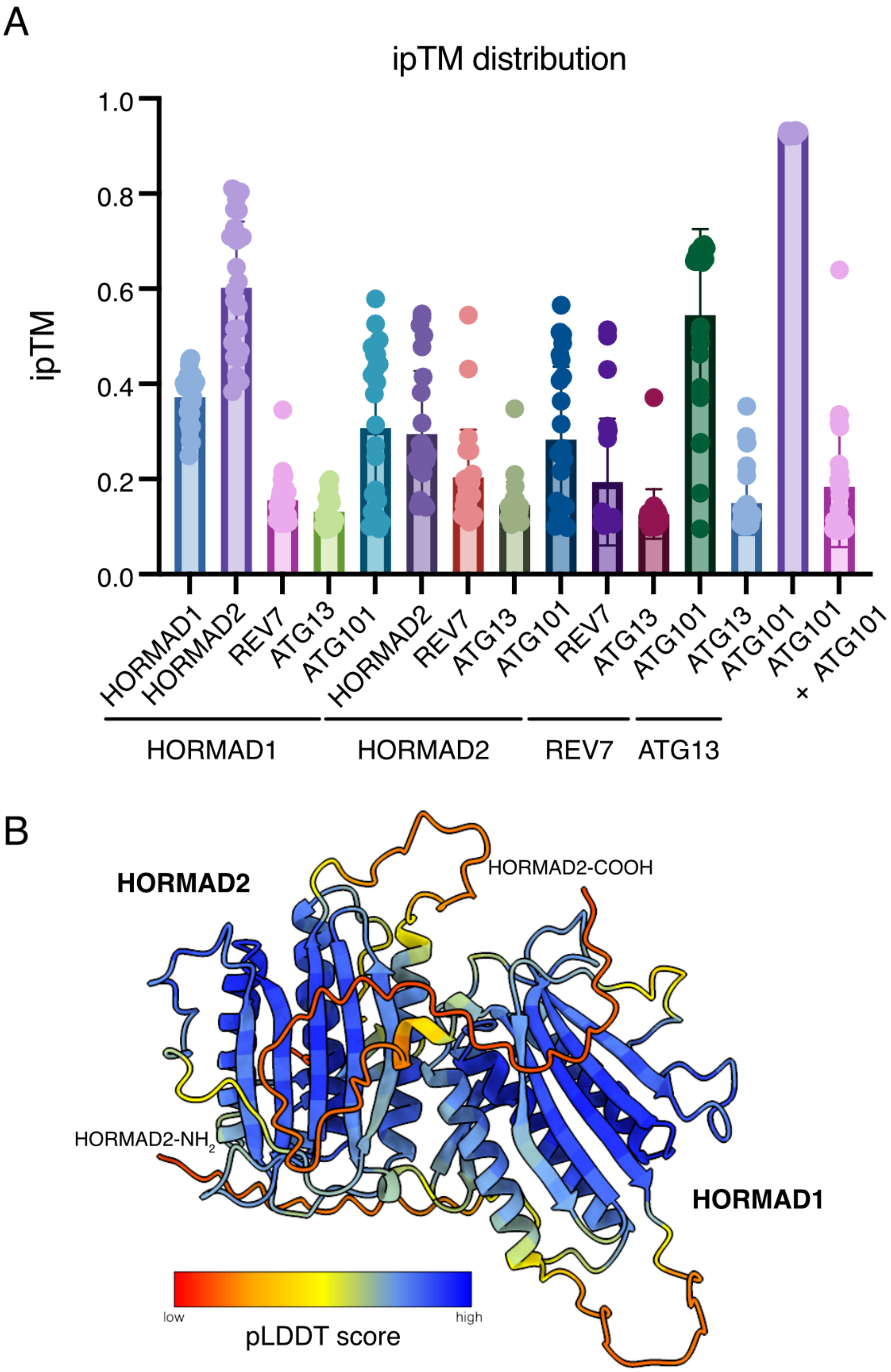


*Supplementary Figure 2 - AlphaFold2 predictions of HORMA dimers*

*A) ipTM distribution of 25 predictions for each indicated combination of HORMA domains. Coloured columns show the mean ipTM value, error bars show standard deviation. B) Top scoring model of HORMAD1$^{HORMA}$ and HORMAD2$^{HORMA}$ (as shown in Figure 2C), coloured by pLDDT score.*

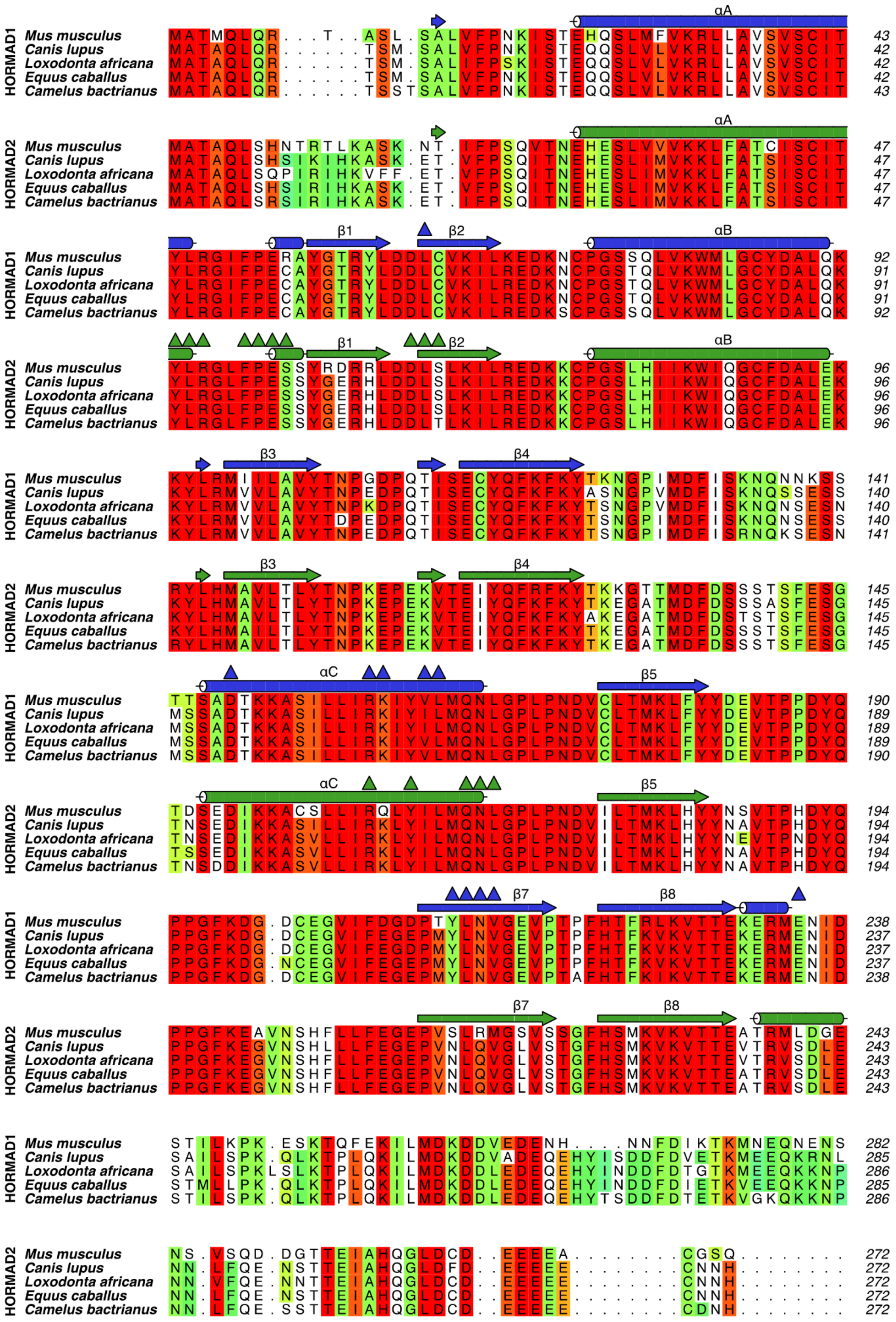
αA
HORMAD1
Mus musculus MATMQLQR...T..ASL.SALVFPNKISTEHQSLMFVKRLLAVSVSCIT 43
Canis lupus MATAQLQR......TSM.SALVFPNKISTEQQSLVLVKRLLAVSVSCIT 42
Loxodonta africana MATAQLQR......TSM.SALIFPSKISTEQQSLVLVKRLLAVSVSCIT 42
Equus caballus MATAQLQR......TSM.SALVFPNKISTEQQSLVLVKRLLAVSVSCIT 42
Camelus bactrianus MATAQLQR......TSSTSALVFPNKISTEQQSLVLVKRLLAVSVSCIT 43
αA
HORMAD2
Mus musculus MATAQLSHNTRTLKASK.NT.IFPSQVTNEHESLVVVKKLFATCISCIT 47
Canis lupus MATAQLSHSIKIHKASK.ET.VFPSQITNEHESLIMVKKLFATSISCIT 47
Loxodonta africana MATAQLSQPIRIHKVFF.ET.VFPSQITNEHESLIMVKKLFATSISCIT 47
Equus caballus MATAQLSHSIRIHKASK.ET.VFPSQITNEHESLIMVKKLFATSISCIT 47
Camelus bactrianus MATAQLSRSIRIHKASK.ET.IFPSQITNEHESLIMVKKLFATSISCIT 47
β1 β2 αB
HORMAD1
Mus musculus YLRGIFPERAYGTRYLDDLCVKILKEDKNCPGSSQLVKWMLGCYDALQK 92
Canis lupus YLRGIFPECAYGTRYLDDLCVKILREDKNCPGSTQLVKWMLGCYDALQK 91
Loxodonta africana YLRGIFPECAYGTRYLDDLCVKILREDKNCPGSTQLVKWMLGCYDALQK 91
Equus caballus YLRGIFPECAYGTRYLDDLCVKILREDKNCPGSTQLVKWMLGCYDALQK 91
Camelus bactrianus YLRGIFPECAYGTRYLDDLCVKILREDKSCPGSSQLVKWMLGCYDALQK 92
β1 β2 αB
HORMAD2
Mus musculus YLRGLFPESSYRDRRLDDLSLKILREDKKCPGSLHIIKWIQGCFDALEK 96
Canis lupus YLRGLFPESSYGERHLDDLSLKILREDKKCPGSLHIIKWIQGCFDALEK 96
Loxodonta africana YLRGLFPESSYGERHLDDLSLKILREDKKCPGSLHIIKWIQGCFDALEK 96
Equus caballus YLRGLFPESSYGERHLDDLSLKILREDKKCPGSLHIIKWIQGCFDALEK 96
Camelus bactrianus YLRGLFPESSYGERHLDDLTLKILREDKKCPGSLHIIKWIQGCFDALEK 96
β3 β4
HORMAD1
Mus musculus KYLRMIILAVYTNPGDPQTISECYQFKFKYTKNGPIMDFISKNQNNKSS 141
Canis lupus KYLRMVVLAVYTNPEDPQTISECYQFKFKYASNGPVMDFISKNQSSESS 140
Loxodonta africana KYLRMVVLAVYTNPKDPQTISECYQFKFKYTSNGPVMDFISKNQNSESN 140
Equus caballus KYLRMVVLAVYTDPEDPQTISECYQFKFKYTSNGPIMDFISKNQNSESS 140
Camelus bactrianus KYLRMVVLAVYTNPEDPQTISECYQFKFKYTSNGPIMDFISRNQKSESN 141
β3 β4
HORMAD2
Mus musculus RYLHMAVLTLYTNPKEPEKVTEIYQFRFKYTKKGTTMDFDSSSTSFESG 145
Canis lupus KYLHMAVLTLYTNPKEPEKVTEIYQFKFKYTKEGATMDFDSSSASFESG 145
Loxodonta africana KYLHMAVLTLYTNPKEPEKVTEIYQFKFKYAKEGATMDFDSTSTSFESG 145
Equus caballus KYLHMAILTLYTNPKEPEKVTEIYQFKFKYTKEGATMDFDSSSTSFESG 145
Camelus bactrianus RYLHMAVLTLYTNPKEPEKVTEIYQFKFKYTKEGATMDFDSSSTSFESG 145
αC β5
HORMAD1
Mus musculus TTSADTKKASILLIRKIYVLMQNLGPLPNDVCLTMKLFYYDEVTPPDYQ 190
Canis lupus MSSADTKKASILLIRKIYILMQNLGPLPNDVCLTMKLFYYDEVTPPDYQ 189
Loxodonta africana MSSADTKKASILLIRKIYILMQNLGPLPNDVCLTMKLFYYDEVTPPDYQ 189
Equus caballus MSSADTKKASILLIRKIYVLMQNLGPLPNDVCLTMKLFYYDEVTPPDYQ 189
Camelus bactrianus MSSADTKKASILLIRKIYVLMQNLGPLPNDVCLTMKLFYYDEVTPPDYQ 190
αC β5
HORMAD2
Mus musculus TDSEDIKKACSLLIRQLYILMQNLGPLPNDVILTMKLHYYNSVTPHDYQ 194
Canis lupus TNSEDIKKASILLIRKLYILMQNLGPLPNDVILTMKLHYYNAVTPHDYQ 194
Loxodonta africana TNSEDIKKASVLLIRKLYILMQNLGPLPNDVILTMKLHYYNEVTPNDYQ 194
Equus caballus TSSEDIKKASVLLIRKLYILMQNLGPLPNDVILTMKLHYYNAVTPHDYQ 194
Camelus bactrianus TNSDDIKKASVLLIRKLYILMQNLGPLPNDVILTMKLHYYNAVTPHDYQ 194
β7 β8
HORMAD1
Mus musculus PPGFKDG.DCEGVIFDGDPTYLNVGEVPTPFHTFRLKVTTEKERMENID 238
Canis lupus PPGFKDG.DCEGVIFEGEPMYLNVGEVPTPFHTFKVKVTTEKERMENID 237
Loxodonta africana PPGFKDG.DCEGVIFEGDPMYLNVGEVPTPFHTFKVKVTTEKERMENID 237
Equus caballus PPGFKDG.NCEGVIFEGEPMYLNVGEVPTPFHTFKVKVTTEKERMENID 237
Camelus bactrianus PPGFKDG.DCEGVIFEGEPMYLNVGEVPTAFHTFKIKVTTEKERMENID 238
β7 β8
HORMAD2
Mus musculus PPGFKEAVNSHFLLFEGEPVSLRMGSVSSGFHSMKVKVTTEATRMLDGE 243
Canis lupus PPGFKEGVNSHLLLFEGEPVNLQVGLVSTGFHSMKVKVTTEVTRVSDLE 243
Loxodonta africana PPGFKEGVNSHFLLFEGEPVNLQVGLVSTGFHSMKVKVTTEVTRVSDLE 243
Equus caballus PPGFKEGVNSHFLLFEGEPVNLQVGLVSTGFHSMKVKVTTEATRVSDLE 243
Camelus bactrianus PPGFKEGVNSHFLLFEGEPVNLQVGLVSTGFHSMKVKVTTEATRVSDLE 243
HORMAD1
Mus musculus STILKPK.ESKTQFEKILMDKDDVEDENH....NNFDIKTKMNEQNENS 282
Canis lupus SAILSPK.QLKTPLQKILMDKDDVADEQEHYISDDFDVETKMEEQKRNL 285
Loxodonta africana SAILSPKLSLKTPLQKILMDKDDLEDEQEHYINDDFDTGTKMEEQKKNP 286
Equus caballus STMLLPK.QLKTPLQKILMDKDDLEDEQEHYINDDFDIETKVEEQKKNP 285
Camelus bactrianus STILSPK.QLKTPLQKILMDKDDLEDEQEHYTSDDFDTETKVGKQKKNP 286
HORMAD2
Mus musculus NS.VSQD.DGTTEIAHQGLDCD..EEEEA........CGSQ......... 272
Canis lupus NN.LFQE.NSTTEIAHQGLDFD..EEEEE........CNNH......... 272
Loxodonta africana NN.VFQE.NNTTEIAHQGLDCD..EEEEE........CNNH......... 272
Equus caballus NN.LFQE.NSTTEIAHQGLDCD..EEEEE........CNNH......... 272
Camelus bactrianus NN.LFQE.SSTTEIAHQGLDCD..EEEEE........CDNH......... 272

*Supplementary Figure 3 - Alignment of HORMAD1$^{HORMA}$ and HORMAD2$^{HORMA}$*

*Secondary structure based on the predicted complex shown. Triangles indicate predicted interaction regions.*

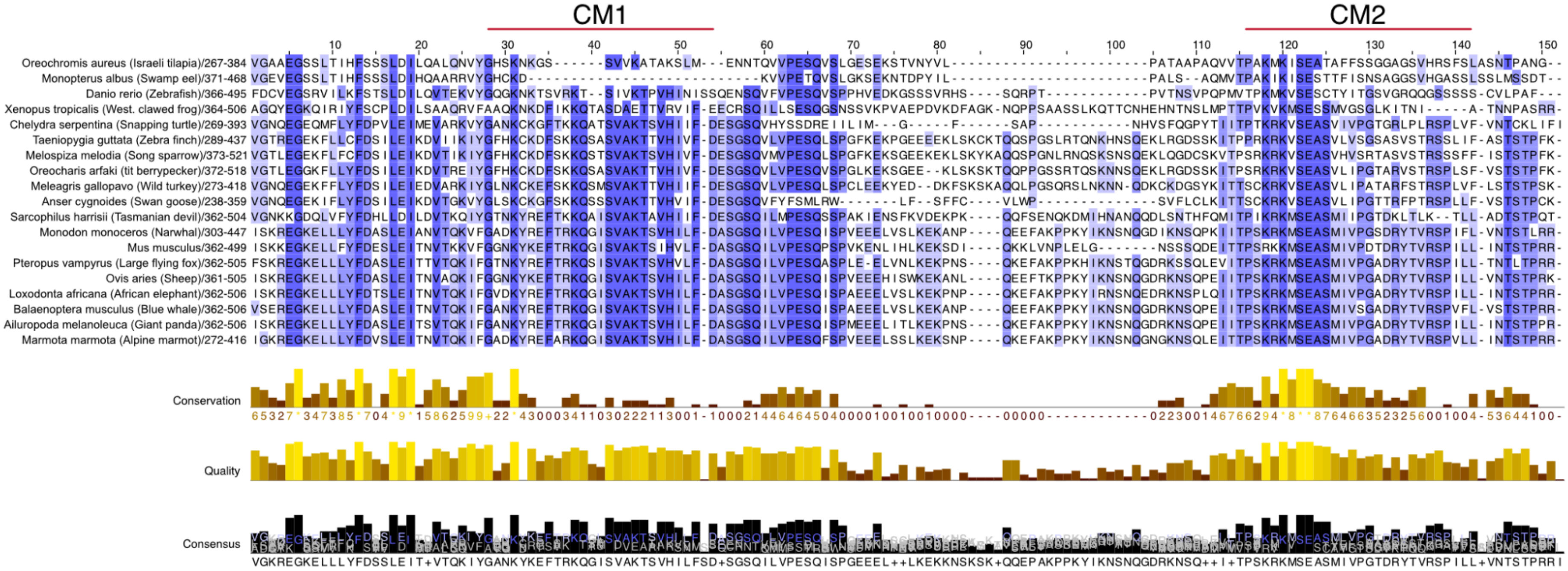

*Supplementary Figure 4 - SYCP2 closure motif region alignments among 19 vertebrates species. Red lines indicate the CM1 (residues in Mus musculus 391-415 (West et al., 2019)) and CM2 (residues in Mus musculus 466-484, this study).*

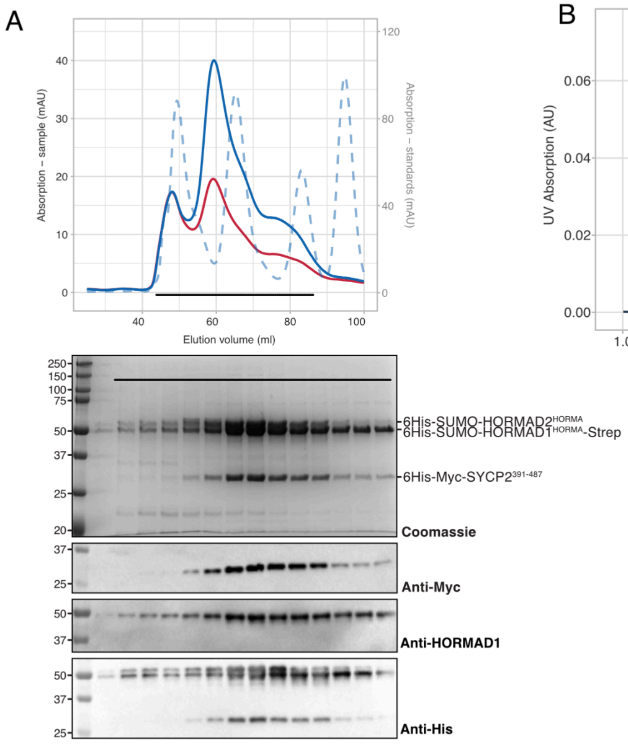


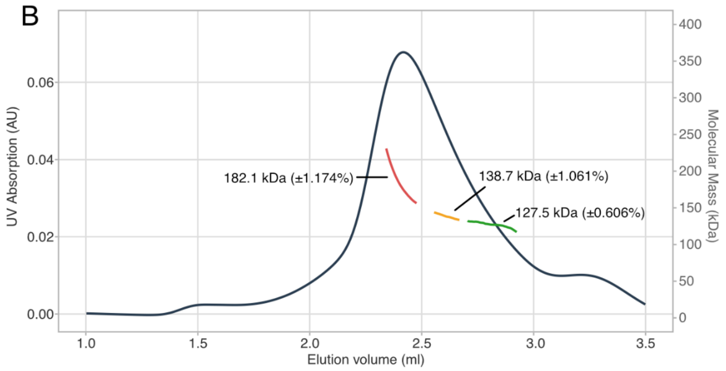


| Myc-SYCP2 $^{391-487}$ 21.84 kDa | SUMO-HORMAD1$^{HORMA}$-Strep 44.24 kDs | SUMO-HORMAD2$^{HORMA}$ 45.24 kDa | kDa |
|---|---|---|---|
| 0 | 1 | 1 | 89.48 |
| 1 | 1 | 0 | 66.08 |
| 1 | 0 | 1 | 67.08 |
| 1 | 1 | 1 | 111.32 |
| 2 | 1 | 1 | 133.16 |
| 2 | 2 | 1 | 177.4 |
| 1 | 2 | 2 | 200 |

*Supplementary Figure 5 - Purification and SEC-MALS analysis of SYCP2$^{391-487}$-HORMAD1-HORMAD2 complex. A. Size-exclusion chromatography of the SYCP2$^{391-487}$-HORMAD1-HORMAD2 complex on the Superdex200 16/600. B. Upper panel: SEC-MALS of SYCP2$^{391-487}$-HORMAD1-HORMAD2 on the column Superdex200 5/150 in 20 mM HEPES pH 8.2, 150 mM NaCl, 1 mM TCEP with the measured molecular masses; lower panel: theoretical masses corresponding to different stoichiometry of potential complex.*
